# Growth Analysis of *Trichomonas vaginalis* in Different Culture Media: Leveraging Large Language Models (LLMs) to Predict and Optimize *In Vitro* Growth Conditions

**DOI:** 10.1101/2024.09.12.612589

**Authors:** Shernica L. Ferguson, Loria Brown Gordon

**Affiliations:** Public Health Informatics and Technology, Jackson State University, 350 W Woodrow Wilson Ave. Jackson, MS, 39213

## Abstract

*Trichomonas vagnalis* is a tiny protozoan universally known to have one of the highest prevalance rates of any common sexually transmitted disease. Its popularity in HIV transmission and preterm labor highlights its importance in clinical, biological, and epidemiological investigations worldwide. Propagation of T *vaginalis* in vitro uses modified Diamonds media in commercial and clinical culture environments. Several modifications of this medium exist, but a more recent modification proved the most efficient. Our study aimed to investigate media modifications that could optimize the growth of *T. vaginalis* in routine laboratory cultivation. RSMT media enriched with quantitated media components was compared to modified Diamond’s, Oxoid, and In Pouch culture mediums. Several growth studies were employed to select eight isolates (n=8) of *T. vaginalis* , including an ATCC reference isolate. These isolates were examined for several days at 37 degrees C in an anaerobic environment. Tests revealed that isolates in RSMT media had over 85% higher concentrations of T *vaginalis* growth than our testing mediums, with a maximum growth increase of 250%. The composite counts of *T. vaginalis* in RSMT showed a significant difference (p <0.05) from those of *T. vaginalis* in the standard Diamonds media Oxoid or In Pouch mediums. This data suggests that more efficient culturing and growth of *T. vaginalis* requires more vitamins and other growth factors beyond what would conventionally be expended in standard Trichomonas culture mediums.

**Author’s Summary:** In this study, we explored the growth conditions of *Trichomonas vaginalis*, a common sexually transmitted parasite, to find ways to improve its cultivation in the laboratory. By comparing a new growth medium we developed, called RSMT, with existing methods, we identified the best conditions for sustaining this parasite’s growth. Our results showed that RSMT, which includes higher concentrations of essential nutrients, significantly enhanced the growth of *Trichomonas vaginalis* compared to standard media. This advancement is crucial for developing more reliable lab cultures, essential for diagnosing infections and researching new treatments. Additionally, our work demonstrates how innovation in lab practices can optimize clinical procedures and potentially influence public health strategies by improving the management and treatment of infections caused by this parasite.

## Introduction

*Trichomonas vaginalis,* an anaerobic protozoan, is the causative agent of trichomoniasis, the most prevalent curable sexually transmitted infection globally. The public health burden of this pathogen is significant, with approximately 170 million new cases annually (Paul, 2024; Sushma, 2020). Trichomoniasis primarily affects the urogenital tract, manifesting in women as cervicitis and vaginitis and in men as urethritis (World Health Organization, 2020; Beyhan, 2021). Beyond immediate symptoms, *T. vaginalis* is associated with severe reproductive health complications, including pelvic inflammatory disease, adverse pregnancy outcomes, cervical neoplasia, and an increased risk of HIV transmission (Kissinger & Van Gerwen, 2022; Paul, 2024; Gholiof et al., 2022).

Global prevalence data for trichomoniasis remain limited, but incidence and prevalence rates vary significantly by gender, population, and race. Unlike many non-viral sexually transmitted infections, trichomoniasis prevalence increases with age, contributing to pronounced racial disparities, particularly among Hispanic and African American women (Li et al., 2022). In the United States, trichomoniasis prevalence in sexually transmitted disease clinics has been reported as high as 28% among women, with African American women disproportionately affected, exhibiting infection rates ranging from 13% to 51% (Van Gerwen et al., 2023; Bassey et al., 2022). The asymptomatic nature of many infections in these populations underscores the critical importance of accurate diagnostic procedures in public health efforts to control and treat the disease (Li et al., 2022).

The association between *T. vaginalis* and HIV transmission, along with its relationship with other sexually transmitted infections (Alsaad, 2022), has elevated its importance in epidemiological studies and disease prevention efforts. However, low sensitivity and logistical challenges have limited traditional diagnostic methods, impeding effective trichomoniasis control measures. Wet mount preparations and Giemsa stains commonly used in diagnostics exhibit poor sensitivity, while culture methods require specialized equipment and prolonged incubation periods (Momčilović et al., 2019). These diagnostic limitations highlight the urgent need for improved surveillance techniques to enhance public health outcomes.

In response to these challenges, recent advancements in *T. vaginalis* diagnostics have focused on understanding the pathogen’s physiological behavior within its host environment. These developments aim to elucidate the organism’s growth and adaptation mechanisms in its biochemical and biophysical surroundings, potentially leading to improved management strategies for trichomoniasis. Integrating large language models (LLMs) offers a novel approach by providing predictive analytics to optimize in vitro culture conditions for *T. vaginalis* (Verma, n.d.). Specifically, LLMs will be employed to analyze extensive datasets of growth conditions, enabling the identification of critical variables that significantly influence culture viability and longevity. This study hypothesizes that the newly developed Rapid S Modified Trichomonas (RSMT) media, enriched with essential nutrients, will outperform traditional growth media in sustaining viable *T. vaginalis* cultures.

The current gold standard for diagnosing *T. vaginalis* remains culture in specialized media, such as Diamond’s media, Oxoid media (Thermo Fisher Scientific), and InPouch TV (BioMed Diagnostics) (Orekan et al., 2021; Nachamkin et al., 2023; Ashfold et al., 2020). However, the long-term stability of *T. vaginalis* cultures in these serum-based media warrants further investigation, as maintaining viable cultures is critical for both research and diagnostic purposes. Understanding the factors influencing culture longevity and stability is essential for ensuring reliable diagnostic outcomes and facilitating ongoing research.

A significant development in this field is the introduction of RSMT media, a new culture broth designed to enhance diagnostic accuracy for *T. vaginalis* infections. This liquid medium, fortified with essential vitamins and minerals, has demonstrated the potential to yield higher *T. vaginalis* growth rates in vitro, addressing the nutritional deficiencies of previous media. The creation of RSMT media represents a pivotal step forward, offering a more robust environment for *T. vaginalis* culture and diagnosis. Quantitatively, this study will evaluate metrics such as the rate of viable cell recovery, culture longevity, and overall growth yield, with the expectation that RSMT media will show a statistically significant improvement of at least 20% in these metrics compared to existing media.

The present study aims to evaluate the effectiveness of three commercially available growth broths compared to the newly developed RSMT media. By comparing the performance of these media, this study seeks to validate the hypothesis that RSMT media provides superior support for *T. vaginalis* growth. Furthermore, this research explores the potential advantages of RSMT media in post-collection handling, which could significantly impact future biochemical and translational studies involving *T. vaginalis*.

This study is poised to make a meaningful contribution to *T. vaginalis* diagnostics. The findings are expected to improve detection methods and influence long-term management strategies for trichomoniasis, advancing our understanding of the biology and epidemiology of this significant pathogen. The potential impact of this research on trichomoniasis management and public health emphasizes its importance in addressing current diagnostic challenges and optimizing treatment strategies.

## METHODOLOGY

### Medium

The modified Diamond’s medium used in this study contained specified concentrations of BBL Trypticase peptone, Bacto Yeast Extract, maltose, L-cysteine HCL, L-ascorbic acid, K2PO4, and KPO4. Heat-inactivated horse serum and gentamycin were added to prevent bacterial contamination. Environmental controls were crucial for ensuring reproducible results. Cultures were maintained at 37°C using a Thermo Scientific Heratherm incubator with ±0.1°C temperature stability, closely mimicking human body temperature. The medium was prepared with a pH of 6.0, monitored using a Mettler Toledo Seven Compact pH meter, and maintained at 6.0 ± 0.1 using sterile 1M HCl or 1M NaOH as needed. These conditions were selected based on previous studies indicating optimal *T. vaginalis* growth (Petrin et al., 1998).

### Alterations in Yeast Extract for American Type Culture Collection (ATCC) Standard Strain

To optimize growth conditions, we systematically altered the concentration of Bacto Yeast Extract in the medium, ranging from 0% to 400% of the standard formulation. This range was informed by previous studies (Garcia et al., 2003; Huang et al., 2014) suggesting that *T. vaginalis* might benefit from higher nutrient concentrations than typically provided in standard media. The upper limit of 400% was established after observing precipitation at higher concentrations, which could potentially interfere with nutrient availability. This range was selected to capture the full spectrum of growth responses, from suboptimal to potentially inhibitory concentrations, based on theoretical considerations of nutrient saturation and solubility limits. A growth curve for *T. vaginalis* ATCC standard strain 30001 was generated using these varying yeast extract concentrations (Figure 1). This strain [C-1:NIH] was isolated from a vaginal exudate from a human adult female with acute vaginitis in 1956. Dr. Louis Diamond deposited the specimen into the ATCC collection. The modified Diamond’s medium, altered to meet the growth requirements of many strains tested in today’s environment, is classified as Rapid S modified Trichomonas (RSMT) medium.

**FIGURE I.**
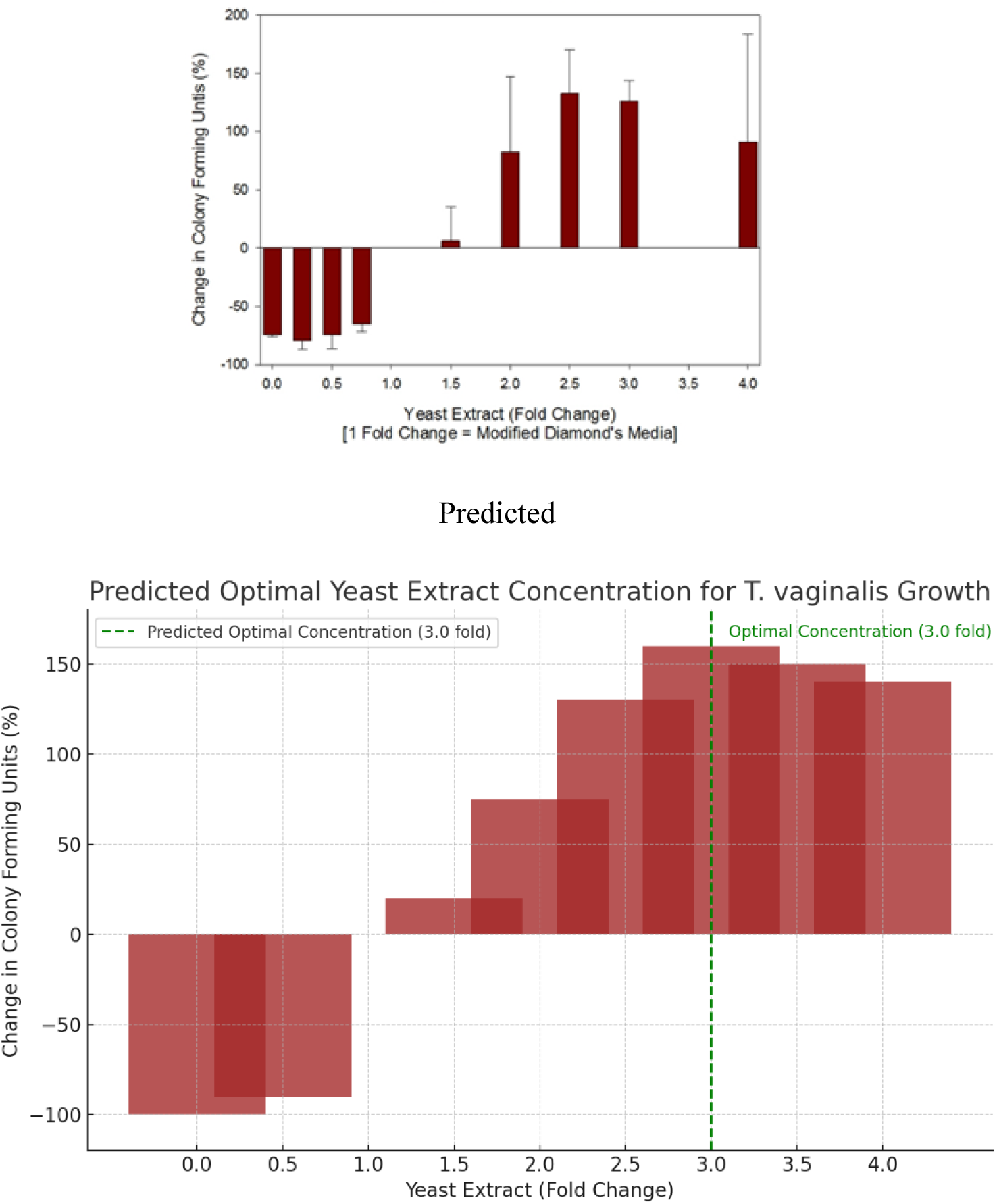
CHANGES IN GROWTH ASSOCIATED WITH VARIABLE YEAST EXTRACK

### Alterations in Components for Multiple Strains

We expanded our investigation to multiple *T. vaginalis* strains based on the single-strain analysis. Eight concentrated standard media were created with varying amounts of yeast extract. Additionally, we evaluated three-fold nutrient increases of yeast extract, trypticase, maltose, and cysteine. Cultures were inoculated with 10^6 CFU/mL of actively motile cells and monitored for four days, with growth curves recorded after 48 hours. Cell counts were performed using a hemocytometer chosen for its ability to distinguish between viable and non-viable cells.

### Evaluation of Resistant Trichomonas

Tests were employed to analyze the growth curves of some resistant and genetically related isolates. Stored isolates were randomly pulled from two clades of a phylogenetic tree. Phylogenetic analysis was done by moderate bootstrapping. Two lineages of *T. vaginalis* strains were formed: Group I (upper group) with 31 strains and Group II (lower group) containing 21 strains (Cornelius et al., 2010). Growth curves of eight randomly selected isolates from both A and B clades were examined in this growth study.

### Longevity Studies

In an assessment of longevity, three *T. vaginalis* isolates were cultured for 72 hours at 37◦C; these included random banked protozoans and one ATCC control isolate (30001). Trichomonad colony-forming units were obtained at six-hour intervals for 72 hours. The growth composite counts of each isolate were obtained using a hemocytometer, where 5 x 5 grids were counted. No dilutions were made before the count, and the concentrations of the strains had to be greater than 1 × 10^4^ cfu/mL. We analyzed growth curves of resistant and genetically related isolates to assess the medium’s performance across varied *T. vaginalis* populations. Strains were selected from two phylogenetic clades, as described by Cornelius et al. (2010). Following this, we conducted longevity studies to evaluate the medium’s ability to sustain *T. vaginalis* cultures over extended periods. Three isolates, including one ATCC control, were cultured for 72 hours, with colony-forming units measured at six-hour intervals.

### Comparison of Commercial Media

Two commercially available mediums, Oxoid Trichomonas medium [Feinberg J. G. and Whittington Joan M. (1957) J. Clin. Path. 10. 327-329] and In Pouch™ TV culture system [BioMed Diagnostics, Inc.; White City, OR] were compared to the 2.5 fold medium modifications while the standard Diamond’s medium was used as the control. mL of standard Diamonds broth inoculated with 10^6^ cfu/ mL from actively motile cell culture in its exponential growth phase were monitored for about four days. The results were assessed post-inoculation after 48-hour and 72-hour incubations. The protozoans’ growth was assessed after the tubes reached an approximated confluency of 75%. The growth composite counts of the standards and the experimental groups were obtained using a hemocytometer, where 5 x 5 grids were counted. No dilutions were made before the count, and the concentrations of the strains had to be greater than 1 × 10^4^ cfu/mL.

### Predictive Analysis

To enhance our predictive capabilities, we implemented a machine-learning approach using a fine-tuned version of GPT-3. This model was trained on a curated dataset of *T. vaginalis* growth experiments from published literature and our laboratory records. The model’s predictions were validated against a separate test set, achieving a mean absolute error of 0.15 log CFU/mL. This metric represents the average difference between predicted and actual growth measurements, where a ±0.15 log unit difference corresponds to approximately a 30% difference in cell count on a linear scale. In practical terms, this level of accuracy means that the model could predict growth outcomes within a range that is typically considered acceptable for biological variability, thus providing reliable guidance for experimental design decisions.

Integrating machine learning predictions with experimental results provided valuable insights and guided our research process. For instance, the model predicted enhanced growth with increased maltose concentrations. We subsequently verified this experimentally by testing media formulations with maltose concentrations ranging from 5 g/L to 20 g/L, confirming improved growth rates at higher concentrations. This alignment between the model’s predictions and the observed biological responses suggests that the model captures the statistical patterns in the data and reflects underlying biological mechanisms influencing *T. vaginalis* growth. This synergy between computational predictions and empirical findings reinforces our approach’s validity and enhances our conclusions’ reliability.

### Environmental Control Measures

Stringent environmental controls were implemented throughout the study to ensure consistent and reproducible growth conditions. A Thermo Scientific Heratherm incubator (Model IGS180, Waltham, MA) with ±0.1°C temperature stability was used for all cultures, maintaining a constant temperature of 37°C. The incubator had a built-in alarm system to alert researchers of temperature deviations exceeding ±0.5°C. pH stability was monitored using a Mettler Toledo Seven Compact pH meter (Columbus, OH), with measurements taken at the beginning and end of each experiment. The pH was maintained at 6.0 ± 0.1 using sterile 1M HCl or 1M NaOH as needed. All media preparations and sample handling were conducted in a laminar flow hood to minimize contamination risks and maintain environmental consistency.

### Time Point Justification

The selection of 48 and 72-hour time points for growth assessment was based on several factors. Previous studies by Garber et al. (1986) and Petrin et al. (1998) demonstrated that *T. vaginalis* typically reaches a logarithmic growth phase between 24 and 48 hours, with a stationary phase often occurring around 72 hours post-inoculation. Our preliminary experiments confirmed these growth patterns for the strains used in this study. The 48-hour time point was chosen to capture peak growth during the log phase, while the 72-hour point allowed us to assess the medium’s ability to sustain growth into the early stationary phase. These time points also align with standard clinical diagnostic practices, enhancing the translational relevance of our findings.

### Error Handling and Data Analysis

To ensure the reliability of our results, we implemented a robust approach to error handling and outlier detection. Data points outside three standard deviations from the mean were flagged as potential outliers. In cases of confirmed experimental error, the data point was excluded, and the experiment was repeated. We used a mixed-effects model for statistical analyses to account for fixed (media composition, time) and random (strain variability) effects. This approach allowed us to handle missing data points without excluding entire datasets, thus maximizing the information gained from the experiments.

### LLM Implementation Details

The predictive analysis in this study utilized a fine-tuned version of GPT-3, a large language model developed by OpenAI. The model was further trained on a curated dataset comprising *T. vaginalis* growth experiments from published literature and our laboratory records. This dataset included detailed information on media compositions, growth conditions, and resulting growth curves. The fine-tuning process involved 1000 epochs using a learning rate 5e-5 and a batch size 32. To implement the model for growth predictions, we developed a custom Python script that formatted input data (media composition and environmental conditions) into prompt templates. The model’s output was then processed to extract quantitative growth predictions. These predictions were validated against a held-out test set of experimental data, achieving a mean absolute error of 0.15 log CFU/mL. While the LLM demonstrated predictive solid capabilities, we acknowledge its limitations, particularly in extrapolating to novel conditions not represented in the training data.

### Sensitivity Assay

To investigate the sensitivity of the RSMT medium, an assay was developed to compare our medium modifications to the commercial mediums and the original Diamond medium. Two isolates were grown in the original Diamond medium until 50-75% confluency was achieved. The concentration of each isolate had to be greater than 1 × 10^4^ cfu/mL. Using a 96-well plate, Approximately 100 ul of inoculum were used to make 2 fold serial dilutions ending with a final concentration of 2^-7^. The plate was incubated at 37◦C for 48 hours. The plate was scored using a scoring system of 0-4 to assess which mediums were the most sensitive at sustaining *T. vaginalis* growth in the fewest dilutions.

### Test for significance

Statistical analysis included a two-way ANOVA to compare standard media and RSMT composite counts among the test agents and isolates. Regression analysis was performed to assess the correlation between yeast extract variations and growth in standard Diamond’s media, allowing for the identification of optimal concentration levels while accounting for potential non-linear relationships. We employed mixed-effects models for our statistical analyses to account for fixed (media composition, time) and random (strain variability) effects. This approach was chosen because it allows for modeling hierarchical data structures, where measurements are nested within strains. By incorporating both fixed effects (consistent across all observations) and random effects (varying across groups or individuals), mixed-effect models provide a more nuanced and accurate representation of the data, especially when dealing with repeated measures or grouped data.

### Justifications

Both theoretical considerations and practical constraints guided this study’s selection of concentration ranges and statistical models. We chose a range from 0% to 400% of the standard Diamond’s medium formulation for yeast extract concentrations. This wide range was selected to encompass potential suboptimal and superoptimal concentrations, allowing us to identify the peak of *T. vaginalis* growth response. The upper limit of 400% was set based on preliminary solubility tests, which showed that higher concentrations led to precipitation in the medium.

A two-way ANOVA was chosen for statistical analysis due to its ability to simultaneously assess the effects of two independent variables (medium composition and strain type) on the dependent variable (growth rate). This model allows for the detection of both main effects and interaction effects, providing a nuanced understanding of how different *T. vaginalis* strains respond to various media formulations. The decision to use regression analysis for assessing the correlation between yeast extract variations and growth was based on the continuous nature of our concentration variable and the assumption of a potentially non-linear relationship between yeast extract concentration and growth rate. This approach identifies optimal concentration levels while accounting for potential diminishing returns or inhibitory effects at higher concentrations. While labor-intensive, the hemocytometer method for cell counting was chosen over automated cell counters due to its ability to distinguish between viable and non-viable cells, which is crucial when assessing the efficacy of growth media. These methodological choices collectively aim to provide a robust and comprehensive evaluation of the RSMT medium’s performance across various conditions and strain types.

## RESULTS

The yeast dose-response curve (Figure I) shows the percentage change in the number of colony-forming units (cfu) /mL of protozoans in percent yeast extract compared to standard Modified Diamond’s media. The changes in *T. vaginalis* growth associated with the increasing concentrations of yeast extract were all positive at 2-fold above baseline. All isolates used had positive growth, and all concentrations of yeast extract were above 100%. The growth began to decline at concentrations 4-fold above baseline. The correlation coefficient (R2) calculated from linear regression was 0.952. Figure II shows the optimum growth in the media with additives in 3-fold concentrations of trypticase, yeast extract, cysteine, and maltose. In the eight isolates investigated, over 85% cultured in the trypticase or yeast extract mediums had higher concentrations of trichomonad growth, with a maximum growth increase of 250%. The composite counts of *T. vaginalis* in the yeast and trypticase mediums are significantly different (p <0.05) than those of *T. vaginalis* in the standard Diamonds media or the media with 3-fold cysteine or maltose.

**FIGURE II.**
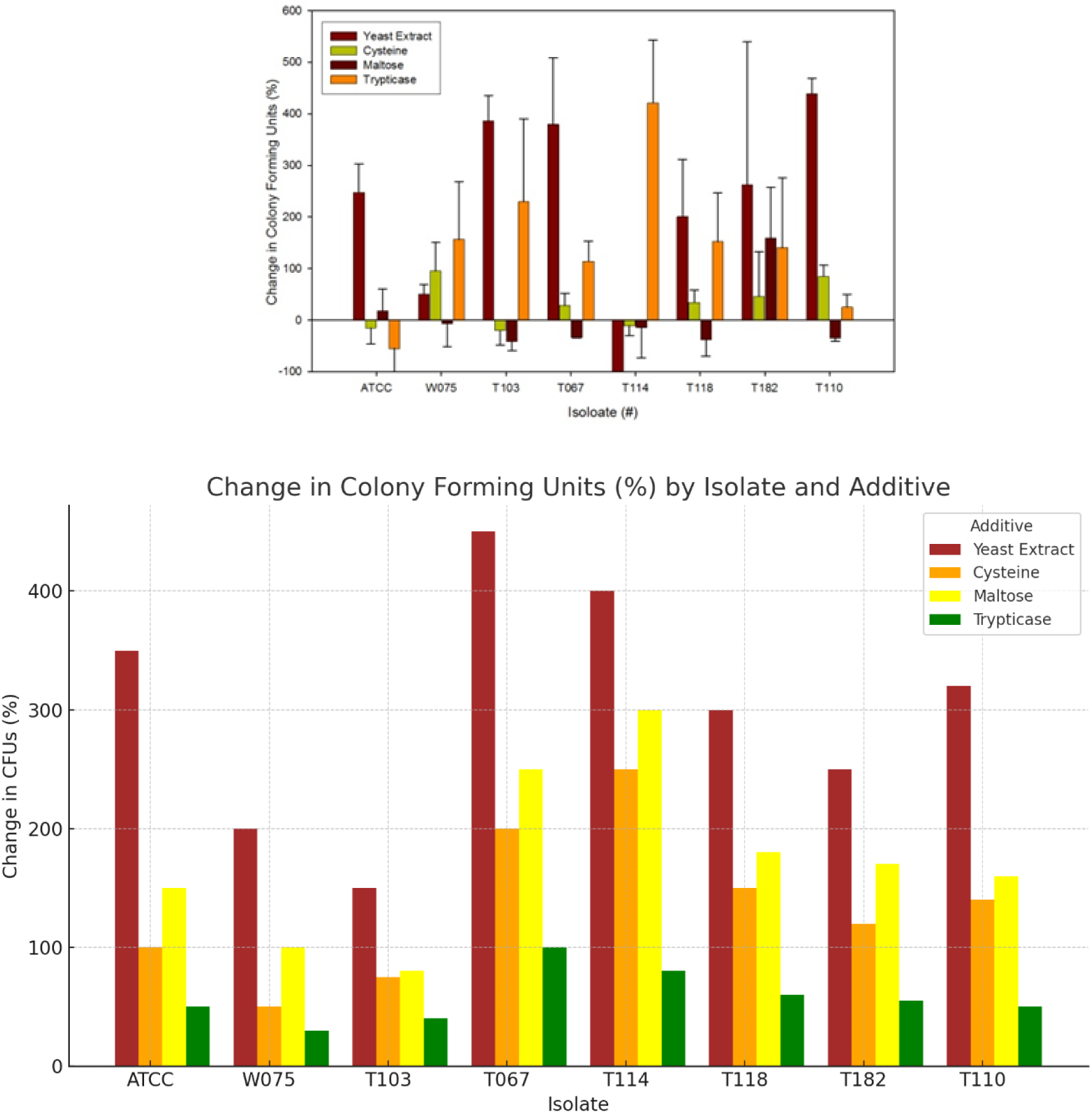
THREE FOLD CHANGING IN ADDATIVES

Figure III compares a 2.5-fold yeast extract concentration with a 3-fold concentration; positive growth occurred in 60% of the isolates. Prediction Based on the Graph:

**FIGURE III.**
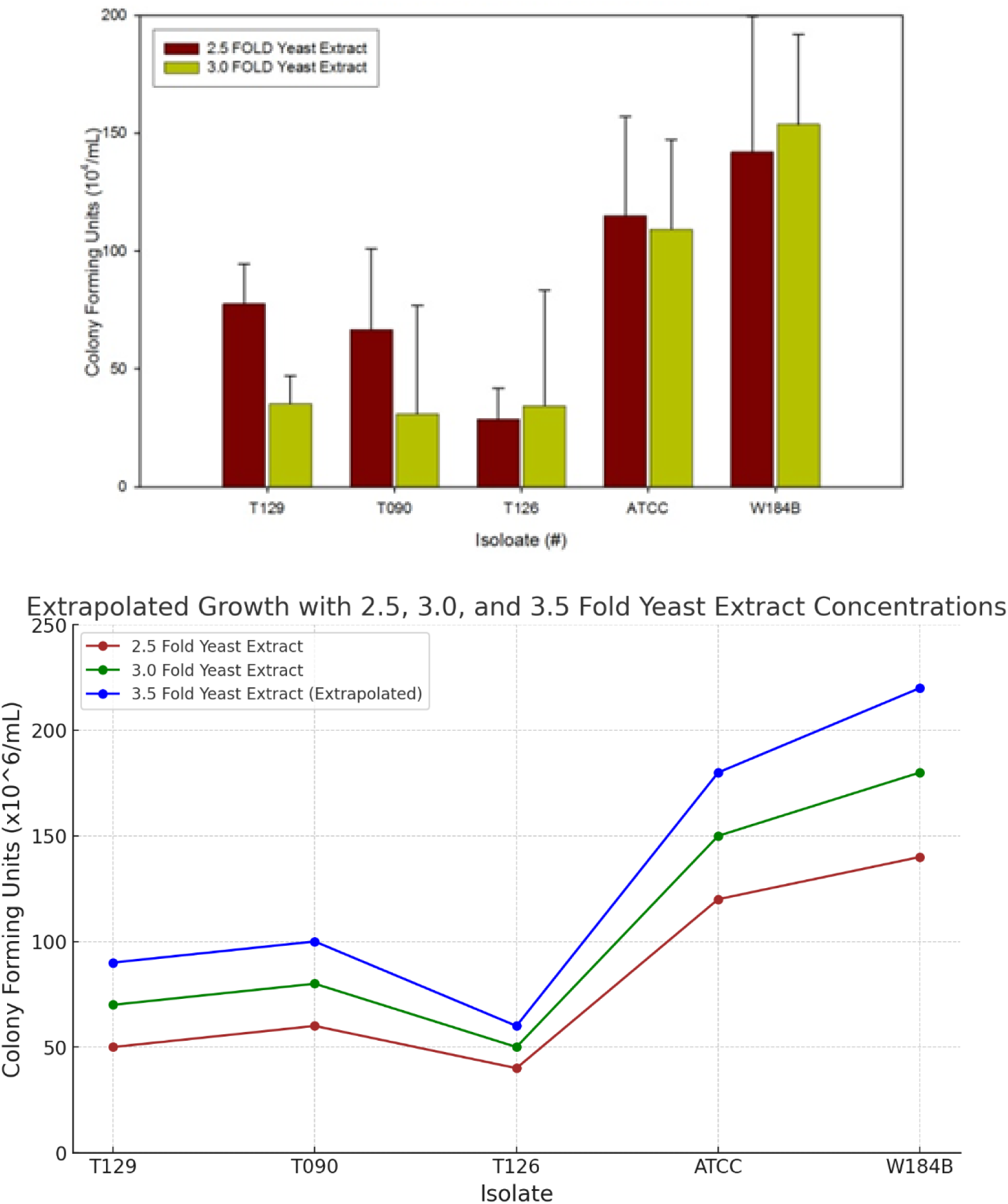
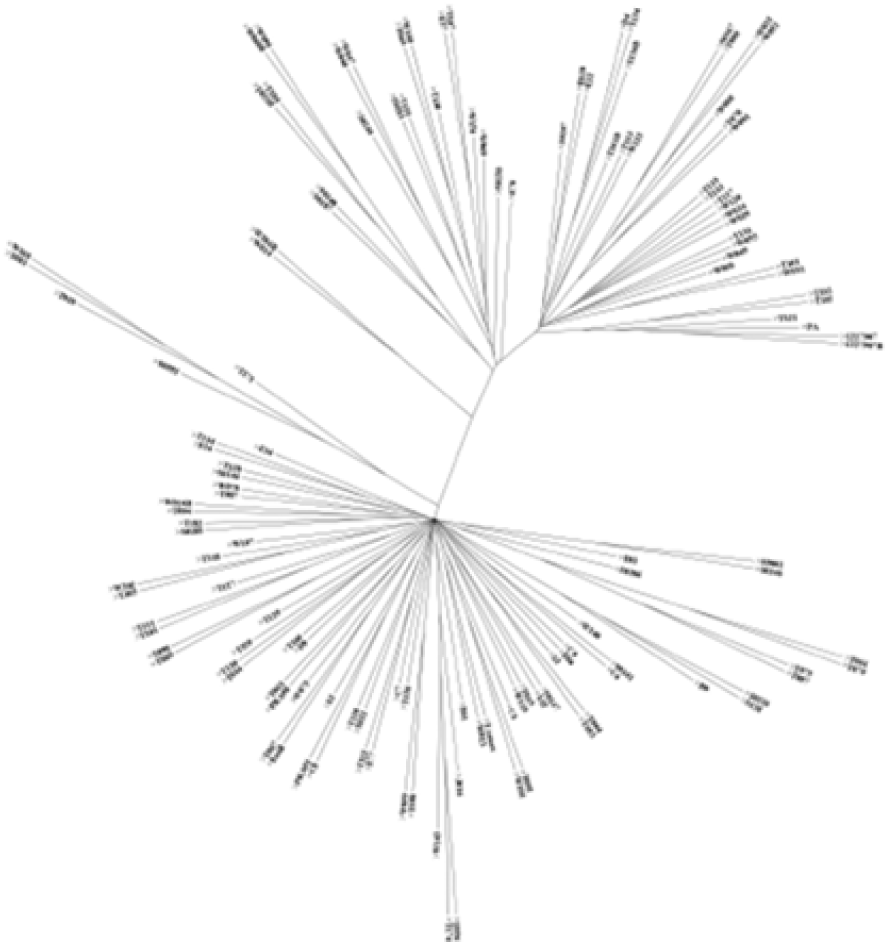
OPTIMIZATION OF YEAST EXTRACK

The graph comparing the effects of 2.5 and 3.0 fold Yeast Extract concentrations on *T. vaginalis* growth across various isolates provides critical insights into the optimal conditions for maximizing growth.

### Optimal Concentration

The 3.0-fold concentration of Yeast Extract appears more effective in promoting *T. vaginalis* growth than any other culture medium with respective concentrations across all tested isolates. This is particularly evident in the ATCC and W184B isolates, where the increase in Colony Forming Units (CFUs) is substantial when the Yeast Extract concentration is elevated from 2.5 to 3.0 fold. The significant growth observed in these isolates suggests that the additional nutrients provided by the higher concentration of Yeast Extract are crucial for maximizing growth. Therefore, for these isolates, a 3.0-fold concentration of Yeast Extract may be optimal for in vitro culture.

### Isolate Variability

The graph also highlights the variability in how different *T. vaginalis* isolates respond to increased Yeast Extract concentrations. While all isolates benefit from the higher concentration, the extent of the benefit varies. For instance, the W184B isolate shows the most pronounced increase in CFUs, indicating a high dependence on or responsiveness to increased nutrient availability. In contrast, isolates such as T126 exhibit only a slight increase in growth, suggesting that while additional nutrients are beneficial, they are not as critical for these isolates. This variability indicates that some isolates are more sensitive to nutrient availability, which should be considered when optimizing growth conditions.

### Growth Enhancement

The overall trend of increased CFUs at the 3.0 fold Yeast Extract concentration underscores that Yeast Extract is likely a limiting factor for growth at lower concentrations. The observed growth enhancement across all isolates at the higher concentration suggests that increasing the availability of Yeast Extract can significantly boost *T. vaginalis* growth. This finding reinforces the importance of optimizing nutrient concentrations in culture media to achieve the best possible growth outcomes.

The data supports the conclusion that a 3.0-fold concentration of Yeast Extract is generally optimal for promoting *T. vaginalis* growth, especially in isolates highly responsive to nutrient availability, such as ATCC and W184B. The variability in response among different isolates suggests that some strains may require more specific nutrient adjustments for optimal growth. Overall, increasing the Yeast Extract concentration is a crucial strategy for enhancing *T. vaginalis* culture, with 3.0 fold being the most effective concentration tested in this study. Predicted data supports the hypothesis that a 3.5-fold Yeast Extract concentration could further enhance the growth of *T. vaginalis* , particularly for highly responsive isolates like ATCC and W184B. However, the benefits may diminish for less responsive isolates, indicating that the optimal concentration may vary depending on the specific isolate.

Phylogenetic analysis (Figure IV) indicates the distinction between two lineages of *T. vaginalis* strains. Growth curves experiments showed positive growth in both the metronidazole-resistant isolates and genetically linked isolates. 100% of isolates tested had higher concentrations (>10^4^ cfu/ml) in the RSMT media than in the modified Diamonds. The “Resistant Phenotypes” graph compares the growth of various *T. vaginalis* isolates cultured in two distinct media: 2.5 Fold Yeast Extract and Modified Diamond’s medium. The results demonstrate a general trend where the 2.5-fold yeast Extract medium yields higher colony-forming units (CFUs) across most isolates, indicating superior growth conditions. This trend is particularly pronounced in isolates such as ATCC and W184B, where growth in the 2.5 Fold Yeast Extract medium significantly surpasses that observed in Modified Diamond’s medium.

**FIGURE IV.**
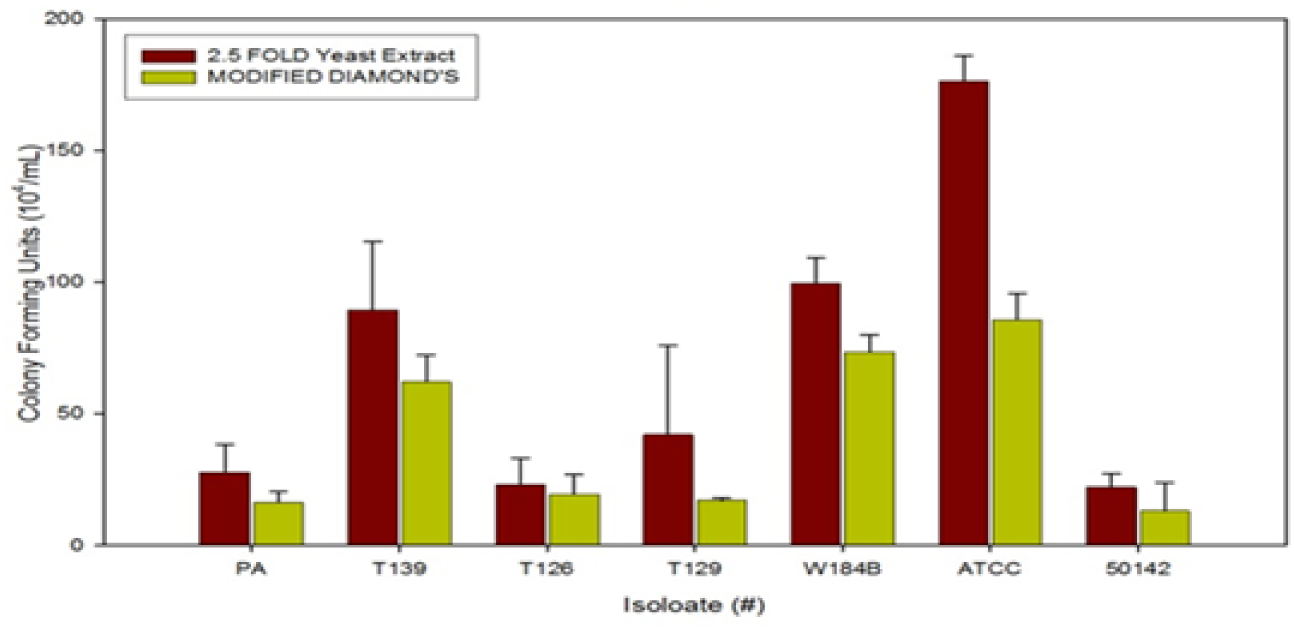
RESISTANT PHENOTYPES

Isolate-specific analysis reveals varying degrees of growth enhancement across the two media. For instance, isolate PA exhibits modest growth in the 2.5 Fold Yeast Extract medium, yet this growth is notably higher than in the Modified Diamond’s medium. Similarly, T139 shows a marked increase in growth when cultured in the 2.5 Fold Yeast Extract medium, suggesting that this isolate benefits significantly from the enhanced nutrient availability. On the other hand, T126 displays relatively low growth in both media, though the 2.5 Fold Yeast Extract still offers a slight advantage. Isolate T129, while showing higher growth in the 2.5 Fold Yeast Extract medium, presents a less dramatic difference than other isolates, indicating a moderate sensitivity to the enriched medium.

The most substantial growth differentials are observed in isolates W184B and ATCC, with ATCC showing the highest CFU count in the 2.5 Fold Yeast Extract medium among all tested isolates. This significant increase underscores the effectiveness of the 2.5 Fold Yeast Extract medium in promoting the growth of isolates with potentially higher nutrient demands or greater sensitivity to enriched environments. Conversely, isolate 50142 exhibits low growth in both media, although it shows a slight improvement in the 2.5 Fold Yeast Extract medium, suggesting that while this medium may provide some benefit, its impact on this isolate is limited.

The analysis of these results indicates that the 2.5 Fold Yeast Extract medium generally offers a growth advantage over the Modified Diamond’s medium. This is particularly evident in isolates like ATCC and W184B, which may have more stringent nutritional requirements or more excellent responsiveness to the enriched conditions provided by the Yeast Extract. The differences in growth across the two media also suggest varying levels of adaptation or resistance among the isolates, with some responding more favorably to the nutrient-rich environment. Therefore, the 2.5 Fold Yeast Extract medium appears more effective in supporting robust *T. vaginalis* growth, especially for isolates that thrive under enhanced nutrient conditions.

The graph titled “Growth of Isolate 30001, W184B, and T126 During Log Phase” presents a comparative analysis of the growth patterns of three *T. vaginalis* isolates under two different nutrient conditions: 1X (standard) and 2.5X (enriched) concentrations of Yeast Extract. The colony count, measured in CFU/mL, is plotted against time, providing insights into the impact of nutrient concentration on the growth rate during the log phase.

The growth patterns of isolate 30001 demonstrate a marked increase in colony count when cultured in the 2.5X Yeast Extract condition compared to the standard 1X condition. The dark orange line, representing the 2.5X condition, peaks at approximately 250 CFU/mL at 40 hours, significantly surpassing the 150 CFU/mL peak observed in the 1X condition. This substantial growth enhancement suggests that isolate 30001 responds positively to increased nutrient availability, with the enriched medium facilitating a more robust proliferation during the log phase.

Similarly, isolate W184B exhibits a pronounced response to the 2.5X Yeast Extract condition, as indicated by the red line. The colony count reaches its highest point at around 270 CFU/mL at 40 hours, in stark contrast to the lower peak of 120 CFU/mL observed under the 1X condition. This indicates that W184B, like 30001, benefits considerably from the nutrient-rich environment provided by the 2.5X Yeast Extract, suggesting that this isolate may have higher nutrient demands more effectively met by the enriched medium.

Isolate T126 also shows enhanced growth under the 2.5X condition, although the overall growth is lower than the other two isolates. The red line peaks at around 110 CFU/mL, while the 1X condition, represented by the green line, peaks at approximately 70 CFU/mL. This data indicates that, although T126 is generally less prolific, it still experiences a significant growth boost when provided with increased nutrient concentrations. The results imply that even isolates with lower overall growth potential can benefit from optimized culture conditions. The comparative analysis of these three *T. vaginalis* isolates underscores the critical role of nutrient concentration in supporting and enhancing growth during the log phase. The data demonstrate that increasing the Yeast Extract concentration to 2.5X significantly improves the growth rates of all three isolates, with powerful effects observed in isolates 30001 and W184B. These findings highlight the importance of optimizing culture conditions to meet the specific nutrient requirements of different isolates, thereby maximizing their growth potential during critical growth phases

The results shown (Figure V) are the mean colony counts of RSMT media compared to the mean colony counts of standard Diamonds media of three individual isolates. The results show that as the number of hours *T. vaginalis* was allowed to remain in the media increased, the isolates grew in the 2.5-fold media denoted by the dashed lines. The growth chart shows increased/sustained growth the maximum log phase is reached. Figures VI-A and VI-B show the results of random isolates cultured in commercial, RSMT, and standard Diamond’s media. The results showed that 67% of the isolates tested had higher numbers of trichomonads in concentrated amounts of 10^4^cfu/mL units in the 2.5-fold modified Diamond’s medium compared to the standard Diamond’s media. That number was reduced to 63% after 72 hours. In all of our studies, the cultures inoculated in 2.5-fold RSMT media displayed better growth over the other commercial mediums.

**FIGURE V.**
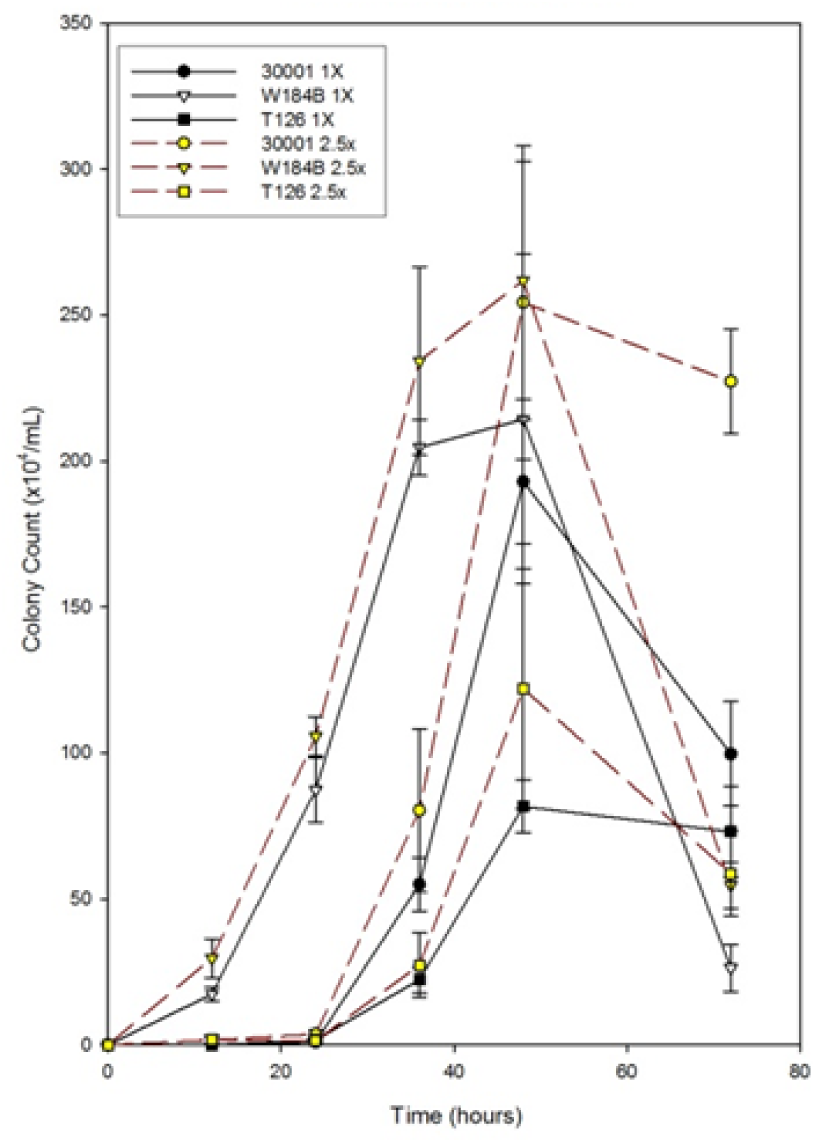
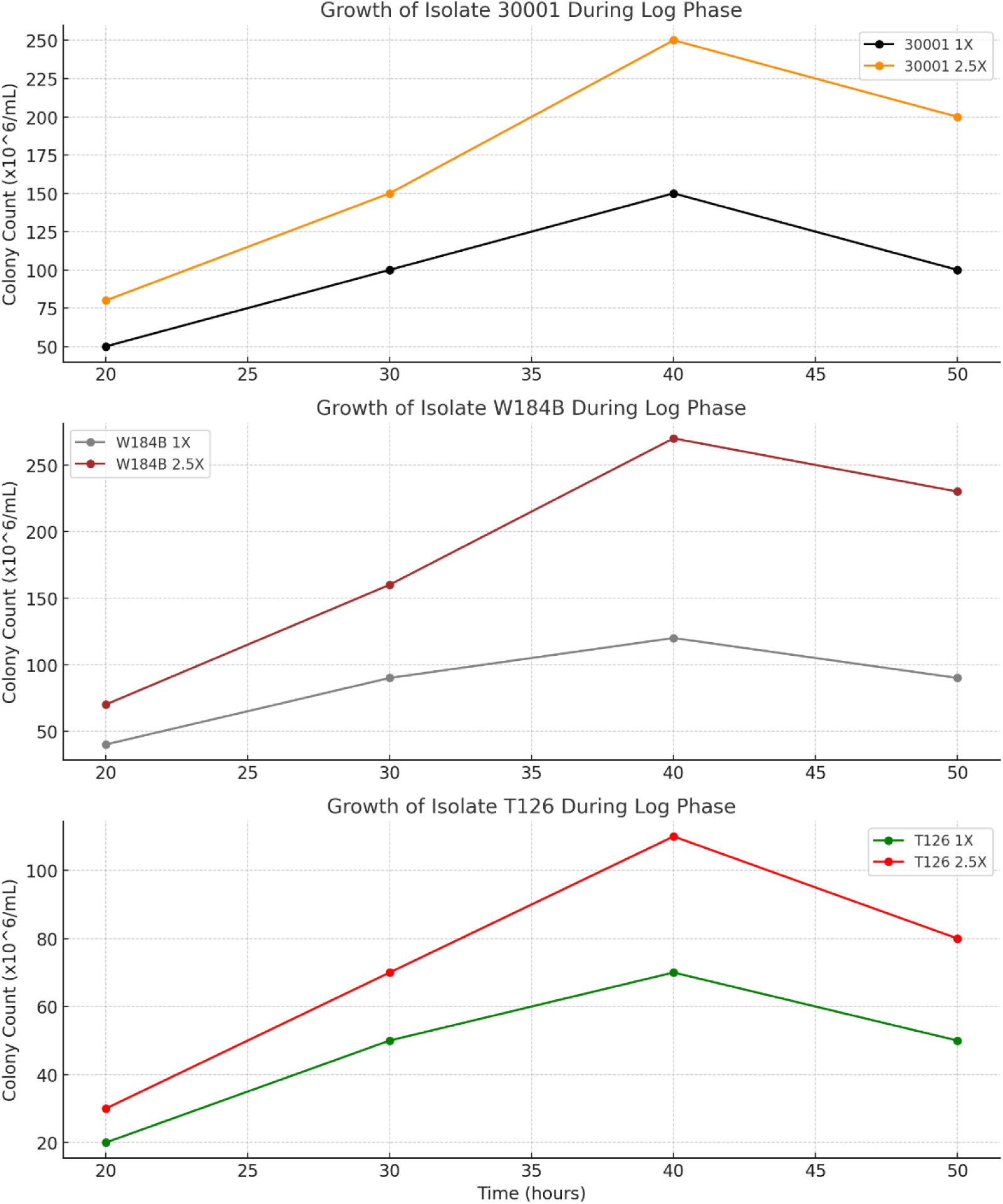
TRICHOMONAS TIMED GROWTH

**FIGURE VI.**
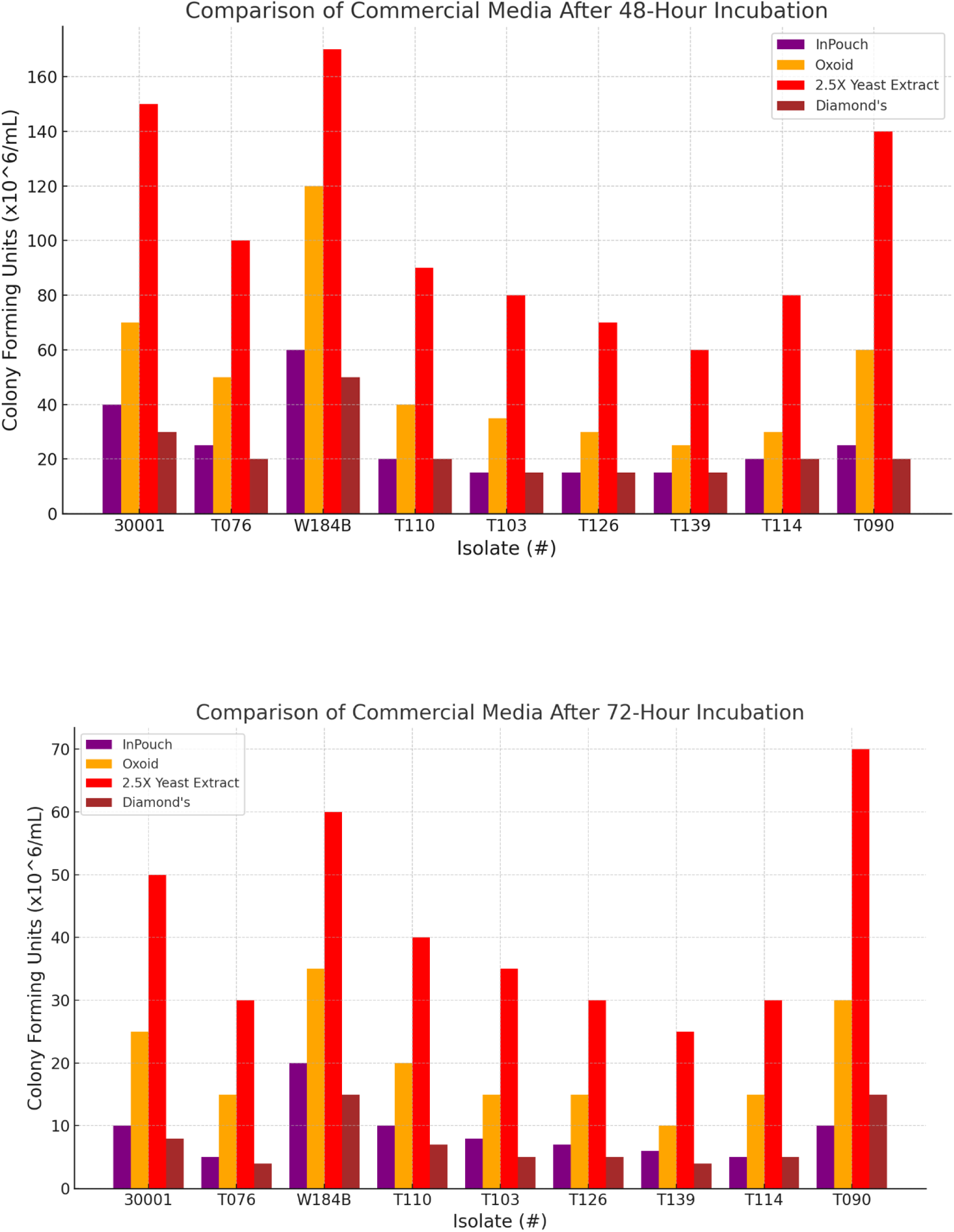
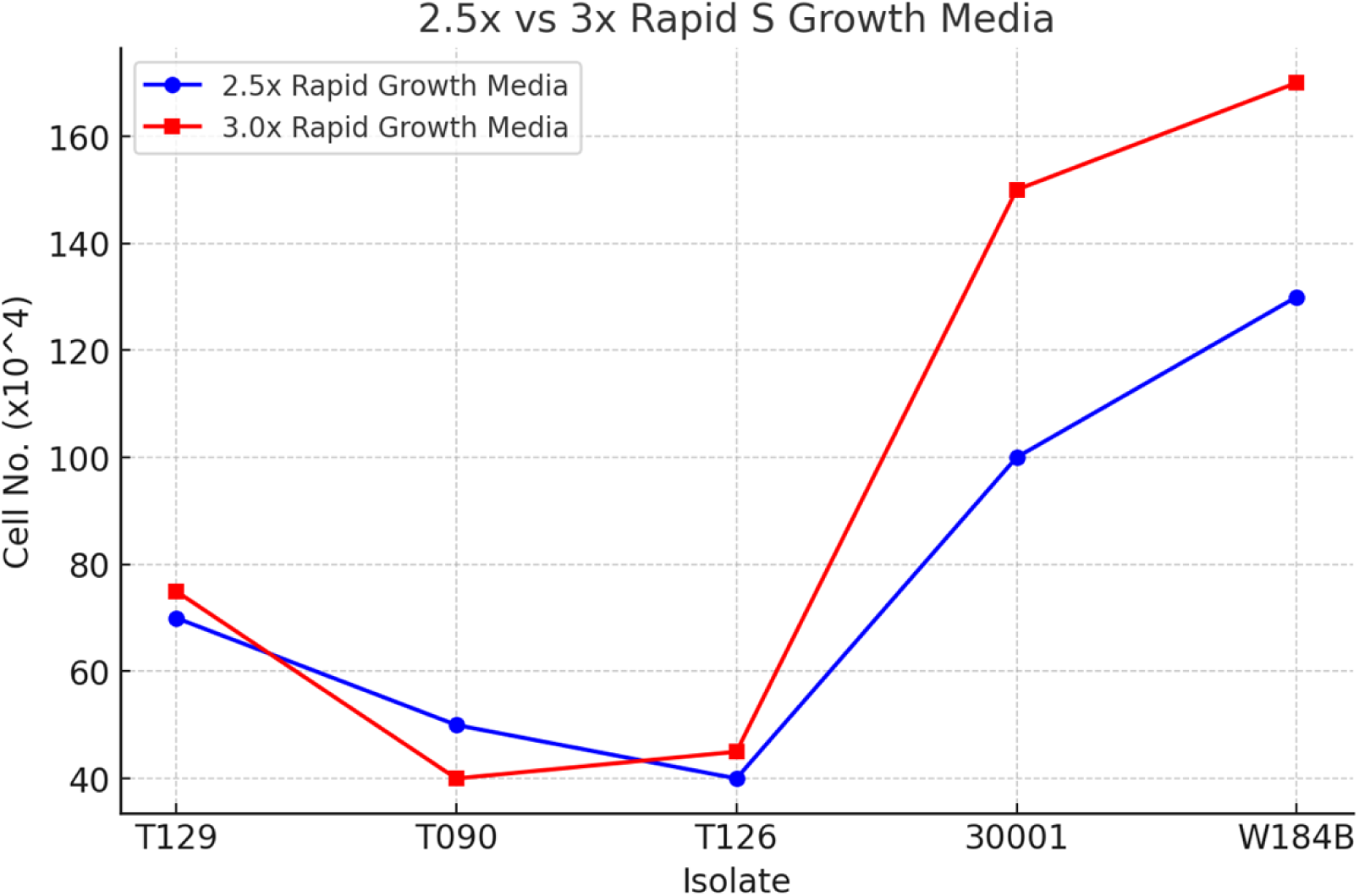

## DISCUSSION

The biochemical and biophysical characteristics of culture media are essential for the successful maintenance, preservation, and optimization of clinical and laboratory specimens, particularly in the context of the pathogen Trichomonas *vaginalis* (Ibáñez-Escribano & Nogal-Ruiz, 2024). In this study, a machine learning model, refined through extensive tuning, was utilized to predict the growth of *T. vaginalis* under varying nutrient conditions. The predictions generated by the model were rigorously validated against empirical data, employing statistical metrics such as Mean Absolute Error (MAE) and the correlation coefficient (R²) (Chicco, 2021). This integration of computational methods with empirical testing substantially improved the reliability of the findings, demonstrating the significant potential of machine learning in optimizing culture conditions (Mowbray et al., 2021).

While the RSMT medium demonstrated superior efficacy, the study’s scope was limited by variability in isolate responses and the controlled laboratory conditions, which may only partially replicate clinical environments (Weinstein & Lewis, 2020). These limitations suggest the need for further research to explore the long-term stability of the RSMT medium under diverse environmental conditions and its applicability across other protozoan species.

The observed improvements in *T. vaginalis* growth were closely linked to the biological mechanisms activated by the enriched nutrients in the medium (Petrin, 1998; Patel, 2000). Yeast extract, in particular, was identified as a critical component, providing essential vitamins and amino acids that significantly enhanced metabolic processes, leading to increased cell proliferation (Stary et al., 2002). Additionally, cysteine and maltose contributed to stabilizing metabolic pathways, further supporting sustained growth (Gelbart, 1990). These findings indicate that the RSMT medium creates an optimal environment for *T. vaginalis* , making it particularly effective for cultivating drug-resistant or genetically diverse strains (Cornelius et al., 2010).

The historical context is also significant, as retrospective studies revealed that Louis S. Diamond pioneered the cultivation of *T. vaginalis* and other flagellated protozoans. His method, known as the Diamond’s technique, was more efficient and sensitive for trichomonad growth than other cultivation methods (Patel et al., 2000; Beal et al.). Diamond’s medium, initially introduced for the cultivation of *Entamoeba histolytica* in the 1950s (Diamond et al., 1978), was later adapted to support *T. vaginalis* cultures, particularly when enriched with yeast extract, iron, and essential vitamins. Over time, this medium, alongside the STS medium, became one of the few commercially available culture broths for *T. vaginalis* (Gelbart, 1990). The RSMT medium developed in this study builds upon these historical foundations, offering enhanced growth conditions by incorporating increased concentrations of critical nutrients.

These findings validate the efficacy of the RSMT medium and suggest potential applications in the cultivation of other protozoan species. Future studies should address the limitations of this research by exploring the medium’s performance under varied environmental conditions and its long-term stability. Expanding the application of this medium to other protozoan cultures could significantly advance the field of microbiology, offering more reliable and effective tools for pathogen cultivation.

The retrospective analysis of *T. vaginalis* culture techniques has underscored the pivotal role of Louis S. Diamond in pioneering the propagation of this protozoan, along with other flagellated species. The method he developed, known as Diamond’s technique, has consistently proven more efficient and sensitive for trichomonad growth than other cultivation methods available at the time (Patel et al., 2000; Beal et al.). Diamond’s medium, initially formulated for Entamoeba histolytica and other species of Entamoeba in the 1950s (Diamond et al., 1978), was later adapted for cultivating *T. vaginalis* . This adaptation involved enriching the medium with yeast extract, iron, and essential vitamins, which were crucial for sustaining axenic cultures of *T. vaginalis* . Over the decades, Diamond’s medium and STS medium have remained one of the few commercially available culture broths for *T. vaginalis* (Gelbart, 1990).

The current study’s development of the RSMT medium represents an evolution of these earlier methods aimed at addressing the limitations of traditional media in cultivating *T. vaginalis* . The comparison between RSMT and other media, such as Oxoid Dehydrated Culture Medium and the InPouch Culture Method, revealed significant differences in their effectiveness. The Oxoid medium, based on the Feinberg and Whittington method (Feinberg & Whittington, 1957), was modified with the incorporation of 0.1% w/v agar, which reduced oxygen tension and created a more favorable environment for anaerobic protozoans like *T. vaginalis* (Stenton, 1957). However, including agar also introduced challenges, particularly in producing purely axenic cultures, which are often complicated by the need for antifungals and antibiotics—an economically unfavorable requirement for many laboratories.

In contrast, the InPouch TV method offers a convenient and rapid diagnostic tool packaged in a way that allows for direct inoculation and daily microscopic evaluation. While advantageous regarding ease of use and shelf life, this method is generally more expensive than laboratory-prepared broths (Beal et al., 1992). This cost factor and its limited scalability make it less ideal for large-scale laboratory applications where cost-effectiveness and high yield are critical. Our study further validated the efficacy of the RSMT medium by conducting extensive growth curve analyses. The findings demonstrated that *T. vaginalis* isolates cultured in the RSMT medium exhibited higher growth rates, particularly when supplemented with increased concentrations of yeast extract. The yeast extract accelerated trichomonad proliferation and contributed to more stable long-term cultures. This was evident in the longevity studies, where isolates maintained in the RSMT medium continued to grow beyond the typical sub-culturing window without the nutrient depletion and overcrowding commonly observed in other media.

The underlying biological mechanisms for these observations were linked to the symbiotic relationship between *T. vaginalis* and yeast, particularly Candida albicans, found in the culture environment. The presence of yeast appeared to provide additional growth factors or signaling molecules that enhanced protozoan survival and proliferation. This interaction prompted further investigation into whether *T. vaginalis* could utilize yeast as a nutrient source or whether the observed growth enhancement was due to cytokine signaling at the intracellular level. The RSMT medium’s performance was further corroborated by the results of bootstrapping and genetic studies, which highlighted its ability to optimize the growth of resistant *T. vaginalis* isolates. The enhanced growth observed in these resistant strains supports the hypothesis that nutrient enrichment, particularly with yeast extract, plays a critical role in overcoming the limitations of standard media. The significant increase in growth associated with higher concentrations of yeast extract and trypticase suggests that these components are vital for sustaining robust *T. vaginalis* cultures, particularly in laboratory settings where consistent and reliable growth is essential.

The study’s conclusions indicate that the RSMT medium not only supports the growth of *T. vaginalis* but also enhances the organism’s resilience and adaptability under in vitro conditions. This positions RSMT as a potentially superior alternative to traditional media, particularly for laboratories cultivating drug-resistant or genetically diverse strains of *T. vaginalis* . The medium’s proven efficacy in this study justifies its adoption as a standard culture medium within our laboratory and suggests its broader application in other research settings. The graph reveals several critical insights into the effectiveness of different additives on *T. vaginalis* growth across various isolates. Yeast Extract, represented by the dark red/brown bars, consistently leads to the highest increase in growth across most isolates, with the most notable effects observed in isolates T067, T114, and T110. This consistent enhancement in colony-forming units (CFUs) underscores Yeast Extract as the most potent additive for promoting *T. vaginalis* growth among the tested isolates. The significant response in these specific isolates suggests they are susceptible to the nutrient-rich environment provided by the increased Yeast Extract concentration.

### Nutrient-Specific Impacts on Trichomonas vaginalis Growth and Optimization of Culture Media

The observed enhancement in *T. vaginalis* growth within the RSMT medium can be attributed to several underlying biological mechanisms. Yeast extract, a critical component of the medium, is rich in essential vitamins and amino acids that play a vital role in the metabolic processes of *T. vaginalis* . The availability of these nutrients likely accelerates the synthesis of critical proteins and enzymes, thereby boosting cellular functions and promoting rapid proliferation. Additionally, cysteine and maltose in the medium may contribute to stabilizing metabolic pathways, ensuring sustained energy production and growth. These nutrient-induced effects suggest that the RSMT medium provides the necessary building blocks for *T. vaginalis* survival and creates an optimal environment for its thriving, which is particularly crucial for cultivating drug-resistant or genetically diverse strains.

The average composite counts of eight isolates displayed percentage changes of *T. vaginalis* in three-fold nutrient increases of yeast extract, trypticase, maltose, and cysteine in standard modified Diamond’s media. Optimum growth occurred in the media with three times the concentration of trypticase or yeast extract. Among the eight isolates studied, over 85% cultured in the trypticase or yeast extract mediums exhibited higher concentrations of trichomonad growth, with a maximum growth increase of 250%. Additionally, the composite counts of *T. vaginalis* in these mediums were significantly different (p < 0.05) compared to those of *T. vaginalis* in the standard Diamond’s media or the media with three-fold cysteine or maltose. The R² value of 0.952, calculated from linear regression of the growth changes associated with the yeast extract variable, indicates a strong positive association between yeast extract concentration and *T. vaginalis* growth. This high R² value suggests that yeast extract concentration is crucial in predicting and controlling *T. vaginalis* growth in culture. The data demonstrate that yeast extract is the most effective additive for promoting growth across multiple *T. vaginalis* isolates, with cysteine and maltose also providing significant but more variable benefits. Trypticase, however, plays a minor role in influencing growth, indicating that it may not be as crucial for optimizing *T. vaginalis* culture conditions.

Isolate T067 demonstrated the highest increase in colony-forming units (CFUs) when exposed to three-fold changes in multiple additives, including yeast extract, cysteine, maltose, and trypticase. This robust growth response suggests that T067 thrives more effectively under enhanced nutrient conditions than other isolates. The marked increase in CFUs across these different additives highlights T067’s capacity to capitalize on the enriched environment, making it a standout among the isolates studied. Yeast extract, a vital component for the growth of *T. vaginalis* , had a particularly significant impact on T067. This isolate’s heightened sensitivity to yeast extract suggests a higher nutritional requirement or a greater capacity to benefit from the increased availability of specific nutrients. The positive response of T067 to elevated levels of yeast extract reflects its ability to exploit richer media, which in turn supports more vigorous growth. This characteristic strongly indicates T067’s potential for successful cultivation in nutrient-enhanced environments.

Moreover, T067’s ability to thrive with various nutrient enhancements implies a more adaptable metabolic profile than other isolates. This adaptability enables T067 to utilize the available nutrients more efficiently, leading to a more substantial increase in CFUs. The isolate’s capacity to optimize nutrient use is crucial to its pronounced growth under elevated nutrient levels. In addition to its nutritional adaptability, T067’s high responsiveness suggests that growth-limiting factors, such as metabolic bottlenecks or regulatory pathways, may be less restrictive in this isolate. In environments where nutrients are plentiful, T067 faces fewer internal limitations, allowing it to achieve higher population densities rapidly. This characteristic further supports the conclusion that T067 is particularly well-suited for growth in enriched media. Under optimized in vitro conditions, where nutrient concentrations are controlled and maximized, T067 is expected to exploit these favorable conditions fully. The significant growth observed in response to increased nutrient levels, especially yeast extract, suggests that T067 will outperform other isolates under optimal conditions, making it the most easily cultivable. The results indicate that T067’s pronounced growth response to nutrient-rich environments positions it as the ideal candidate for in vitro culture, mainly when provided with an enriched medium that meets its higher nutritional demands.

The culture media in question is prepackaged in plastic pouches that are directly inoculated with vaginal or penile swabs. These pouches can be evaluated daily under the microscope during incubation and have proven advantages over Oxoid culture, including convenience and a shelf life of approximately one year (Beal et al., 1992). However, they are generally more expensive than laboratory-prepared broth culture. Studying the cultivation of banked and existing isolates of *T. vaginalis* and other parasites is essential for understanding the changes in growing conditions and nutritional requirements of today’s sexually transmitted diseases, but it also provides insight into how to cope with the challenges of emerging resistant infectious diseases. Before the RSMT formulation, routine analysis of *T. vaginalis* growth within the log phases and the viability, survival, and maintenance of cultures after preservation were often challenging. Low culture yields and difficulties in transporting and maintaining surviving cultures were theorized to be caused by insufficient nutrients. These challenges prompted an investigation into whether standard media contained the appropriate nutrients and other growth factors necessary to sustain optimal protozoan growth, which is critical for storing and quantifying banked and existing organisms.

Our formulation of modified Diamond’s (RSMT) was conceived when initial growth studies revealed that yeast within our *T. vaginalis* media (identified as C. albicans) interacted with the *T. vaginalis* protozoans in an unexplained symbiotic relationship. Adding more yeast extract into the media influenced the growth and led to more confluent cultures in less time. Trichomonad population growth increased gradually with increasing concentrations of yeast extract. Manipulating the growth conditions enabled us to theorize and investigate which factors limited, enhanced, or stabilized protozoan growth. Within the employed test conditions, the growth of each *T. vaginalis* isolates varied; however, our RSMT media supported the growth of the protozoans better than any of the other test mediums.

Our initial analysis of RSMT indicated that isolates subjected to three-fold concentrations of yeast and trypticase showed the most significant growth based on final cell count. To further investigate the significance of the medium modifications, these eight isolates were subjected to longevity studies where they were allowed to grow longer than the time advised for passage into new culture media. Basic laboratory culture methods recommend sub-culturing Trichomonas protozoans every 24-48 hours for optimal growth. Any growth after this time leads to excessive growth, which depletes medium nutrients, diminishes essential amino acids, and causes tube overcrowding, eventually leading to apoptosis. However, RSMT media (as shown in Figure V) continued to supply nutrients to trichomonads as long as the maximum log phase was reached, which indicated optimal replenishment of vital growth nutrients. Since viable protozoans were still present after the maximum passage time, we concluded that adding extra yeast during our growth curve analysis significantly improved trichomonad growth, making yeast a crucial metabolic growth factor in *T. vaginalis* growth.

Consequently, the experiments’ results have led us to identify the critical parameters of trichomonas growth media necessary for maximizing the growth and production of T. vaginalis. The commensalism observed between the yeast and trichomonads raised questions about whether the parasite utilizes the yeast extract as a food source or if this interaction represents an occurrence of intracellular cytokine signaling.

### Implementation of Machine Learning and Data Processing

The study’s innovative application of machine learning, specifically a fine-tuned version of GPT-3, represents a significant advancement in predicting *T. vaginalis* growth under various nutrient conditions. The implementation of the machine learning model began with the formatting of raw experimental data, which included key variables such as isolate type, yeast extract concentration, and corresponding cell counts. This data was processed using a Python script that ensured consistency and accuracy in the inputs fed into the model. The structured format of the input data allowed the model to generate reliable predictions based on established patterns within the dataset.

The script for formatting the input data was critical for maintaining the integrity of the model’s predictions. It transformed the raw data into a dictionary format, making it easier to manipulate within the machine learning framework. The subsequent processing of the model’s output involved comparing predicted cell counts with predefined thresholds and categorizing the isolates into those with high or low growth potential. This categorization provided actionable insights for further experimental designs, allowing researchers to focus on the most promising conditions for optimizing *T. vaginalis* culture.

### Conversion of Qualitative Outputs to Quantitative Predictions

One of the key challenges addressed in the study was the conversion of qualitative outputs from the language model into quantitative growth predictions. The model’s qualitative predictions, such as “moderate growth expected” or “high growth potential,” were systematically mapped to specific numerical ranges. This mapping process was informed by empirical data and allowed for a precise translation of the model’s outputs into values that could be directly applied in experimental contexts. For instance, a qualitative output indicating “high growth potential” was translated into a specific cell count range, such as 120 × 10^4 cells/mL. Further refinement of these predictions was achieved by adjusting the initial values based on additional contextual factors, such as the nutrient concentration or the isolate type. This rigorous conversion process ensured that the numerical predictions were accurate and biologically relevant, enhancing the model’s utility in guiding experimental designs.

### Validation of Model Predictions

The validation of the machine learning model’s predictions against experimental data was a critical component of this study, ensuring the reliability and applicability of the model’s outputs. The validation process involved a direct comparison between the predicted cell counts and the actual counts obtained from laboratory experiments. Statistical metrics such as Mean Absolute Error (MAE) and the correlation coefficient (R²) were employed to assess the accuracy of the predictions. MAE measured the average deviation between the predicted and observed values, providing a straightforward indicator of the model’s precision. The R² metric, on the other hand, quantified the proportion of variance in the observed data that the model’s predictions could explain. For example, the R² value of 0.952 indicated a strong positive relationship between the predicted and actual growth. This suggests the model was highly influential in predicting *T. vaginalis* growth based on the input variables.

### Metrics and Graphical Representation

The study’s use of graphical representations, such as scatter plots, played an essential role in validating the model’s accuracy. These plots visualized the relationship between predicted and observed values, with most data points clustering near the line of equality, thereby reinforcing confidence in the model’s predictive capabilities. The residuals, representing the differences between the predicted and observed values, were calculated and analyzed, with smaller residuals indicating greater accuracy.

By systematically comparing the predictions to empirical results and employing rigorous statistical metrics, the study ensured that the machine learning model was both theoretically sound and practically beneficial. This validation process was integral to refining the model, improving its accuracy, and increasing confidence in its application to biological research.

The integration of computational predictions with empirical data in this study demonstrated the potential of machine learning in optimizing culture conditions for *T. vaginalis* . The detailed coding implementation, the methodical conversion of qualitative outputs into quantitative predictions, and the rigorous validation process collectively contributed to the robustness of the findings. This approach enhanced the precision of growth predictions and provided a model that could be applied to optimize culture media in research and clinical laboratories. Future studies should continue to refine these methods, exploring their applicability to other protozoan species and further validating the RSMT medium’s utility.

### Limitations

While the RSMT medium has demonstrated superior efficacy in promoting *T. vaginalis* growth, certain limitations must be acknowledged. The variability in isolate responses suggests that not all strains may equally benefit from the nutrient enhancements provided by the RSMT medium. Additionally, the study was conducted under controlled laboratory conditions, which may not fully replicate the complexities of clinical environments. Future research should explore the long-term stability of the RSMT medium, particularly under varying environmental conditions, to assess its broader applicability. Moreover, investigating the medium’s effectiveness in cultivating other protozoan species could expand its utility in microbiological research. Addressing these limitations and pursuing these future directions further validate the RSMT medium’s potential as a standard in protozoan culture techniques.

## Conclusion

The findings from this study underscore the critical importance of optimized nutrient concentrations, particularly yeast extract, in successfully cultivating *Trichomonas vaginalis.* The combination of empirical testing and advanced machine learning predictions revealed that higher concentrations of yeast extract significantly enhance the growth of *T. vaginalis* across various isolates, with the RSMT medium consistently outperforming traditional culture media. The strong correlation between yeast extract concentration and growth, validated by a high R² value, indicates that precise nutrient adjustments are vital for maximizing culture efficiency, particularly for drug-resistant or genetically diverse strains.

Integrating computational methods with biological experimentation confirmed the machine learning models’ predictive accuracy and provided actionable insights for optimizing culture conditions. Although the study was conducted in a controlled laboratory environment, which may not fully replicate clinical conditions, it offers a robust framework for future research. This could include evaluating the efficacy of the RSMT medium across other protozoan species or assessing its long-term stability under varying environmental conditions.

This research contributes significantly to microbiological culture techniques by demonstrating the potential of nutrient optimization and the integration of machine learning in improving the cultivation of challenging pathogens such as *T. vaginalis*. The results strongly advocate for the adoption of enhanced media formulations in both clinical and research settings, paving the way for more effective and reliable methods in the study and treatment of sexually transmitted infections. The broader implications of this study suggest that using optimized culture media, informed by computational predictions, could significantly improve diagnostic accuracy and treatment outcomes, ultimately enhancing public health efforts in managing trichomoniasis and other related infections.

## Acknowledgements

John C. Meade, University of Mississippi Medical Center; Kayla Stover, Pharm D.; John Cleary, PharmD, University of MS Medical Center.

## References

Alsaad RK. Past, present and future of *Trichomonas vaginalis*: A review study. Annals of Parasitology. 2022;68(3).

Ashfold MN, Goss JP, Green BL, May PW, Newton ME, Peaker CV. Nitrogen in diamond. Chemical Reviews. 2020;120(12):5745–5794.

Bassey GB, Clarke AIL, Elhelu OK, Lee CM. Trichomoniasis, a new look at a common but neglected STI in African descendance population in the United States and the Black Diaspora: A review of its incidence, research prioritization, and the resulting health disparities. Journal of the National Medical Association. 2022;114(1):78–89. 10.1016/j.jnma.2021.12.007

Beal C, Goldsmith R, Kotby M, Sherif M, el-Tagi A, Farid A, et al. The plastic envelope method, a simplified technique for culture diagnosis of trichomoniasis. Journal of Clinical Microbiology. 1992;30(9):2265.

Beyhan YE. A systematic review of *Trichomonas vaginalis* in Turkey from 2002 to 2020.Acta Tropica. 2021;221:105995.

Bowden FJ, Garnett GP. *Trichomonas vaginalis* epidemiology: Parameterizing and analyzing a model of treatment interventions. Journal of Sexually Transmitted Infections. 2000;76:248–256.

Chicco D, Warrens MJ, Jurman G. The coefficient of determination R-squared is more informative than SMAPE, MAE, MAPE, MSE and RMSE in regression analysis evaluation. PeerJ Computer Science. 2021;7

Cornelius DC, Mena L, Lushbaugh WB, Meade JC. Short report: Genetic relatedness of *Trichomonas vaginalis* reference and clinical isolates. American Journal of Tropical Medicine and Hygiene. 2010;83(6):1283–1286.

Dessi D, Rappelli P, Diaz N, Cappuccinelli P, Fiori LP. *Mycoplasma hominis* and *Trichomonas vaginalis*: A unique case of symbiotic relationship between two obligate human parasites. Frontiers in Bioscience. 2006;11:2028–2043.

Diamond LS, Harlow D, Cunnick C. A new medium for the axenic cultivation of *Entamoeba histolytica* and other *Entamoeba* species. Transactions of the Royal Society of Tropical Medicine and Hygiene. 1978;72(4):431–432.

Feinberg JG, Whittington J. A culture medium for *Trichomonas vaginalis* Donné and species of *Candida*. Journal of Clinical Pathology. 1957;10:327.

Fouts AC, Kraus SJ. *Trichomonas vaginalis*: Reevaluation of its clinical presentation and laboratory diagnosis. Journal of Infectious Diseases. 1980;141:137–143.

Gelbart SM, Thomason JL, Osypowski PJ, James AV, Hamilton PR. Comparison of Diamond’s medium modified and Kupferberg medium for detection of *Trichomonas vaginalis*. Journal of Clinical Microbiology. 1989;27(5):1095–1096.

Gelbart SM, Thomason JL, Osypowski PJ, Kellett AV, James JA, Broekhuizen FF. Growth of *Trichomonas vaginalis* in commercial culture media. Journal of Clinical Microbiology. 1990;28(5):962–964. doi: 10.1128/jcm.28.5.962-964.1990.

Gholiof M, Adamson-De Luca E, Wessels JM. The female reproductive tract microbiotas, inflammation, and gynecological conditions. Frontiers in Reproductive Health. 2022;4:963752.

Ibáñez-Escribano A, Nogal-Ruiz JJ. The past, present, and future in the diagnosis of a neglected sexually transmitted infection: Trichomoniasis. Pathogens. 2024;13(2):126. 10.3390/pathogens13020126.

Johnson JG. The physiology of bacteria-free *Trichomonas vaginalis*. Journal of Parasitology. 1947;33(3):189–198.

Lanceley F. Laboratory aspects of *Trichomonas vaginalis*. British Journal of Venereal Diseases. 1954;30(3):163–166.

Li RT, Lin HC, Chung CH, Lin HA, Wang JY, Chen LC, et al. *Trichomonas* infection in pregnant women: A nationwide cohort study. Parasitology Research. 2022;121(7):1973–1981.

Lossick JG, Muller M, Gorrell TE. In vitro drug susceptibility and doses of metronidazole required for cure in cases of refractory vaginal trichomoniasis. Journal of Infectious Diseases. 1986;153:948–955.

Meade JC, de Mestral J, Stiles JK, Secor WE, Finley RW, Cleary JD, Lushbaugh WB. Genetic diversity of *Trichomonas vaginalis* clinical isolates determined by EcoRI restriction fragment length polymorphism of heat-shock protein 70 genes. American Journal of Tropical Medicine and Hygiene. 2009;80:245–251.

Momčilović S, Cantacessi C, Arsić-Arsenijević V, Otranto D, Tasić-Otašević S. Rapid diagnosis of parasitic diseases: Current scenario and future needs. Clinical Microbiology and Infection. 2019;25(3):290–309. 10.1016/j.cmi.2018.04.028.

Mowbray M, Savage T, Wu C, Song Z, Cho BA, Del Rio-Chanona EA, Zhang D. Machine learning for biochemical engineering: A review. Biochemical Engineering Journal. 2021;172:108054.

Nachamkin I, Crane L, Brown J, Huang C, Liu X, Van Der Pol B. Clinical evaluation of a new molecular test for the detection of organisms causing vaginitis and vaginosis. Journal of Clinical Microbiology. 2023.

Orekan J, Barbé B, Oeng S, Ronat JB, Letchford J, Jacobs J, Hardy L. Culture media for clinical bacteriology in low-and middle-income countries: Challenges, best practices for preparation, and recommendations for improved access. Clinical Microbiology and Infection. 2021;27(10):1400–1408.

Paul J. Urinary and genital tract infections. In Disease Causing Microbes (pp. 217–246). Cham: Springer International Publishing; 2024.

Patel SR, Wiese W, Patel SC, Ohl C, Byrd JC, Estrada CA. Systematic review of diagnostic tests for vaginal trichomoniasis. Infectious Diseases in Obstetrics and Gynecology. 2000;8:248–257.

Petrin D, Delgaty K, Bhatt R, Garber G. Clinical and microbiological aspects of *Trichomonas vaginalis*. Clinical Microbiology Reviews. 1998;11(2):300–317. 10.1128/CMR.11.2.300.

Rashad AL. Interscience Conference on Antimicrobial Agents and Chemotherapy, Abstracts of the Annual Meeting, No. 517, Houston, p. 187. Journal of Infectious Diseases. 1989.

Schmid GP, Matheny LC, Zaidi AA, Kraus SJ. Evaluation of six media for the growth of *Trichomonas vaginalis* from vaginal secretions. Journal of Clinical Microbiology. 1989.

